# Identification of mCMQ069, a novel antimalarial with potential for a single-dose cure and/or 28-day chemoprevention

**DOI:** 10.1101/2024.07.01.601566

**Authors:** Anil K. Gupta, James Pedroarena, Armen Nazarian, Jenya Antonova-Koch, Frank Weiss, Krittikorn Kumpornsin, Victor Chi, Ashley K. Woods, Kyoung-Jin Lee, Sean B. Joseph, Shuangwei Li, Jason Brittain, Eugenio De Hostos, James Duffy, Anne Cooper, Peter G. Schultz, Case W. McNamara, Arnab K. Chatterjee

## Abstract

In efforts towards eliminating malaria, a discovery program was initiated to identify a novel antimalarial using KAF156 as a starting point. Following the most recent TCP/TPP guidelines, we have identified mCMQ069 with a predicted single oral dose for treatment (∼40-106 mg) and one-month chemoprevention (∼96-216 mg). We have improved unbound MPC and predicted human clearance by 18-fold and 10-fold respectively when compared to KAF156.

## INTRODUCTION

Malaria continues to be a devasting disease within endemic regions of Africa and Southeast Asia,^1^ and two pockets of locally acquired malaria have recently been identified in the United States.^2^ According to the WHO Malaria Report 2023, there were more than 249 million cases of malarial infection worldwide, leading to ∼608,000 deaths in 2022.^1^ Children are one of the most vulnerable because they generally lack immunoprotection that is acquired from prior, and repeated, malaria infections.^3^ Mosquirix is a relatively new WHO recommended vaccine developed via a collaboration between researchers at GlaxoSmithKline Biologics and Walter Reed Army Institute of Research that is being administered to the youngest children in hopes of offering increased protection.^4^ To date, the efficacy in children under 2 years of age is moderate (< 40%) so additional interventions are necessary to further prevent and control malaria infections.^5^ WHO recommended R21/Matrix-M vaccine in 2023 and it shows good efficacy: 75% when a 3-dose regimen is given seasonally, and 66% when given on an agebased schedule.6 However, these vaccines are only partially protective and, therefore, must be deployed in addition to chemoprevention and other interventional strategies and treatments.

The effective deployment of seasonal malaria chemoprevention (SMC) is one such notable intervention due to its high impact. This strategy has been used in the Sahel region of Africa and, in 2021 alone, more than 45 million children were treated.^7^ This strategy involves oral dosing of an antimalarial regimen at monthly intervals for a maximum of four months. The administration of SMC regimens corresponds with the rainy season (typically 4 months in duration) because this is when malaria transmission is highest. The repeated, monthly dosing aims to treat any existing infections to block parasite transmission back to the mosquito vector and to sustain efficacious plasma exposures in healthy individuals that may become subsequently infected.

The three-drug regimen for Seasonal malaria chemoprevention (SMC comprises sulfadoxine-pyrimethamine (SP) and amodiaquine (AQ) and is also known as SPAQ. The target population are children between 3 and 59 months of age. Concerns about SMC include rebound of the disease due to loss of immunity at older ages, unwanted interruptions in campaigns, and development of drug resistance. Hence, SMC is not recommended for areas where high levels of resistance to SP or AQ have been demonstrated. This includes the seasonal transmission belt in southern Africa where SP resistance is well documented. Additionally, the rise of artemisinin resistance is a big danger to malaria treatment and elimination.^8^ As a result, there is an urgent need to identify novel antimalarials to combat the new emerging resistance to known drugs. Traditional dosing occurs for three days to ensure >>99.9% reduction of parasitemia enabling cure from malaria. In order to improve adherence and effectiveness, there has been a shift in focus towards finding a single-dose cure. Currently, M5717 (Merck), artefenomel (Sanofi) and SJ-733 (Eisai) are the lead programs in Phase 2 trials with a potential single-dose cure, although SJ-733 + cobicistat is being tested in a Phase 2 trial with a 3-day dosing regimen.^9^ Artefenomel is being tested with multiple partner drugs including a novel drug, presumably to identify an effective single-dose combination. There are an additional 3 and 6 candidates in Phase 1 and preclinical development stage respectively with a predicted single-dose or low-dose treatments.^10^ While promising, many of them have either high cost of good (COG) or potential for resistance, providing a unique opportunity for new antimalarials with low COGs and cost of treatment (COT). Furthermore, M5717 is the only oral candidate projected to provide a potential for chemoprophylaxis.

KAF156/Ganaplacide (Novartis) is the most advanced novel antimalarial candidate in development entering a Phase 3 clinical trial in 2024 as a three-day treatment. In a Phase 2 study, a single dose of 800 mg KAF156 alone achieved a 67% 28-day cure rate (NCT01753323).^11^ A follow-up Phase 2 trial of two doses of KAF156 (400 mg/day) in combination with the partner drug lumefantrine (960 mg/day) achieved a 91–98% 28-day cure rate (NCT03167242).^11, 12^ A subsequent drug-drug interaction (DDI) Phase 1 study (NCT05236530) suggests that the Phase 3 study is likely to be a 3-day treatment of KAF156 (400 mg/day) with lumefantrine (960 mg/ day).^13^ The requirement of daily dosing up to 2–3 days for ∼100% 28-day cure rate will lead to high cost of treatment and increased pill burden. Uniquely, KAF156 is pan-active against the parasite lifecycle in the vertebrate host with a novel mode of action, and the first non-ACT in more than 40 years. On the clearance front, recently disclosed data of *in vivo* rat metabolism from Novartis reveals its met-ID profile including detection of polar metabolites. While most of the polar metabolites originated from metabolism of the piperidine ring, hydrolytic cleavage led to the generation of anilino ketoamide. We hypothesized that modification of the 2-position of the imidazole ring of the template could lead to the reduced formation of oxalamide-based metabolite and improved overall clearance without significantly changing overall physiochemical properties. Pan-activity along with underexplored structure-activity-relationship (SAR) work in certain regions of the molecule provide an excellent opportunity to optimize KAF156 further to develop a potentially, best-in-class single-dose treatment and/or 28-day prophylaxis.

Herein, we report mCMQ069, a next-generation KAF156, with improved unbound minimum parasiticidal concentration (MPC) by ∼18x fold and predicted human clearance by 10x fold as compared to KAF156, thereby providing an opportunity for a single-dose cure and chemoprevention of malaria.

## RESULTS

### Structure-Activity Relationship studies

Given the relatively limited SAR published in 2-aryl (Ar) region of the molecule, we undertook a simple modification strategy of that ring. As outlined in Table 1, substitution of the pair of position with halogens or other substituents has a very steep effect on anti-parasitic activity including Table 1 Entry 3 the 4-bromo analog as well as other electron donating and withdrawing groups (Table 1 Entries 4-6). The only tolerated substituent is in Table 1 Entry 7 with substitution with the small alkyl groups such as methyl, so we also undertook some meta- and para-substitution patterns and saw no significant improvement. However, we were very interested in Table 1 Entry 12 with 3,4-difluoro which appeared to be beneficial in terms of potency albeit by about two-fold relative to selectivity.

**Table 1.**
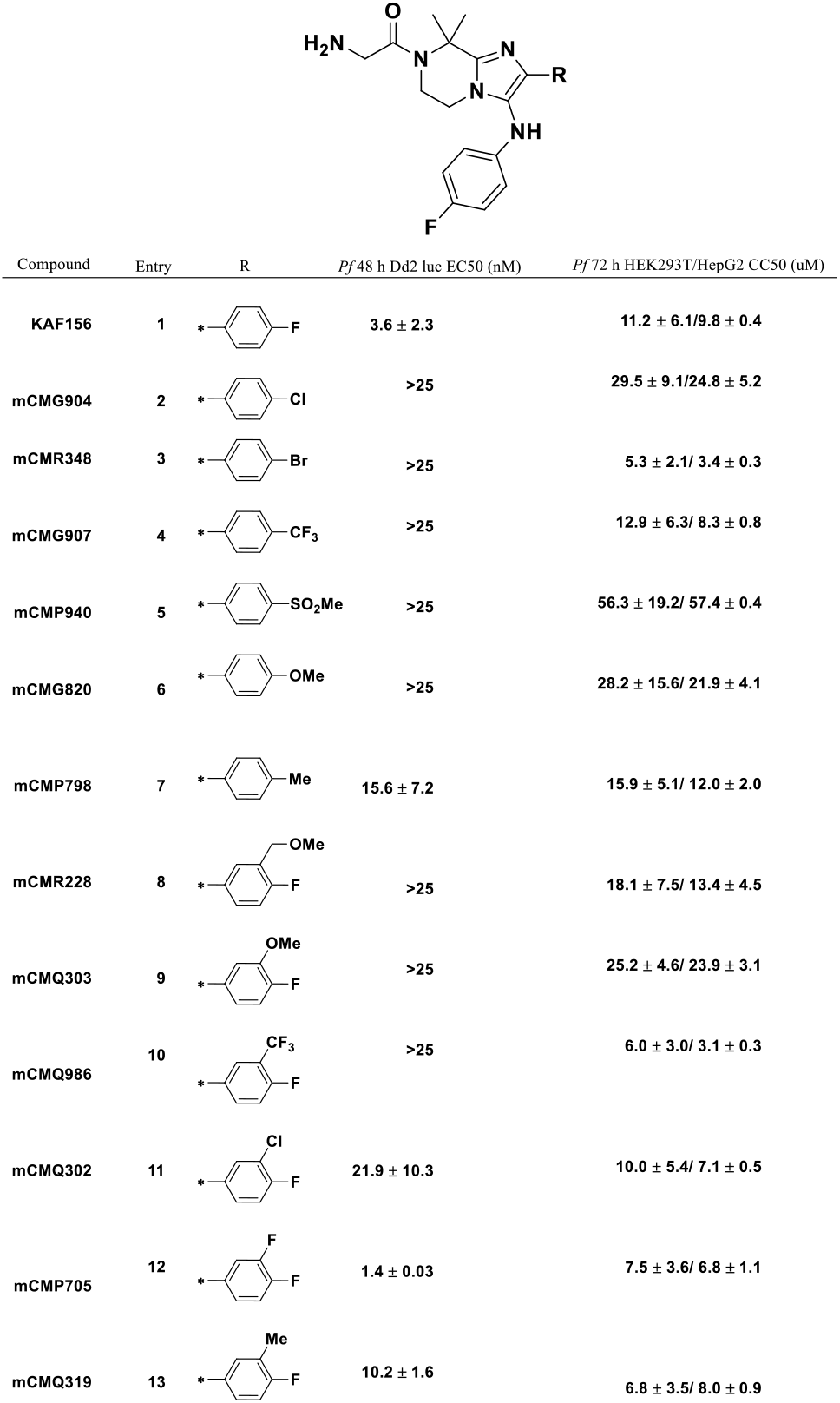
Initial SAR for 2-Ar ring.

| Compound | Entry | R | Pf 48 h Dd2 luc EC50 (nM) | Pf 72 h HEK293T/HepG2 CC50 (uM) |
| --- | --- | --- | --- | --- |
| KAF156 | 1 |  | 3.6 ± 2.3 | 11.2 ± 6.1/9.8 ± 0.4 |
| mCMG904 | 2 |  | >25 | 29.5 ± 9.1/24.8 ± 5.2 |
| mCMR348 | 3 |  | >25 | 5.3 ± 2.1/ 3.4 ± 0.3 |
| mCMG907 | 4 |  | >25 | 12.9 ± 6.3/ 8.3 ± 0.8 |
| mCMP940 | 5 |  | >25 | 56.3 ± 19.2/ 57.4 ± 0.4 |
| mCMG820 | 6 |  | >25 | 28.2 ± 15.6/ 21.9 ± 4.1 |
| mCMP798 | 7 |  | 15.6 ± 7.2 | 15.9 ± 5.1/ 12.0 ± 2.0 |
| mCMR228 | 8 |  | >25 | 18.1 ± 7.5/ 13.4 ± 4.5 |
| mCMQ303 | 9 |  | >25 | 25.2 ± 4.6/ 23.9 ± 3.1 |
| mCMQ986 | 10 |  | >25 | 6.0 ± 3.0/ 3.1 ± 0.3 |
| mCMQ302 | 11 |  | 21.9 ± 10.3 | 10.0 ± 5.4/ 7.1 ± 0.5 |
| mCMP705 | 12 |  | 1.4 ± 0.03 | 7.5 ± 3.6/ 6.8 ± 1.1 |
| mCMQ319 | 13 |  | 10.2 ± 1.6 | 6.8 ± 3.5/ 8.0 ± 0.9 |

Given the result with 3,4-difluoro substitution, we became interested in further probing other regioisomers of difluoro as well as perfluoro analogs that are outlined in Table 2. The position of the substitutions is critical to drive potency in the Dd2 Plasmodium assay. Fluorine substitution is amenable at the meta-position in Table 2 Entry 2 as there is a drop in potency when the fluorine that is moved to the ortho-carbon. Systematic placement of di- and tri-fluoro groups are outlined in Table 2 with several promising compounds emerging with good potency related to KAF156 including Table 2 Entry 4 among others. We concluded that additional fluorines would not reduce potency. We then decided to scan several anilines keeping mCMP705 as a control. Unfortunately, we did not observe any significant improvements in potency in any of the new analogs when compared to mCMP705 (Table 3). Analogs with comparable potency resulted in higher intrinsic clearance (>7 uL/min per million cells) than mCMP705 (Table 3, entries 2,3 and 7). Thereafter, we were keen to get key profiling done in more definitive in vitro Plasmodium assay along with ADME/PK assays for selected analogs as shown in Table 4. Several lead compounds including KAF156 as a positive control were tested in the definitive hypoxanthine NF54 Plasmodium assay along with in vitro human hepatic clearance and rat/mouse in vivo clearance (Table 4). That is where we observed that the 3,4-trifluoro compound Table 4 Entry 9 had 4-fold improved potency relative to KAF156 with similar human hepatocyte clearance and improved rat/mice in vivo clearance. Often these Plasmodium assays are highly correlative, but there can be exceptions to this and so upon repeat we observed that potency improvement was conserved.

**Table 2.** Scanning of multiple fluorines in 2-Ar ring.

| Compound | Entry | R | *Pf* 48 h Dd2 luc EC50 (nM) | *Pf* 72 h HEK293T/HepG2 CC50 (uM) |
| --- | --- | --- | --- | --- |
| KAF156 | 1 |  | 3.6 ± 2.3 | 11.2 ± 6.1/9.8 ± 0.4 |
| mCMN583 | 2 |  | 2.9 ± 0.07 | 9.8 ± 4.0/9.5 ± 1.7 |
| mCMQ022 | 3 |  | 10.7 ± 1.4 | 47.8 ± 24.5/32.9 ± 0.4 |
| mCMP705 | 4 |  | 1.4 ± 0.03 | 7.5 ± 3.6/ 6.8 ± 1.1 |
| mCMP706 | 5 |  | 4.5 ± 0.5 | 23.7 ± 12.7/ 16.4 ± 2.1 |
| mCMR395 | 6 |  | 5.6 ± 1.8 | 17.4 ± 7.2/ 14.4 ± 1.0 |
| mCMR144 | 7 |  | 5.9 ± 1.4 | 18.8 ± 8.3/ 10.0 ± 2.7 |
| mCMQ291 | 8 |  | 4.5 ± 0.5 | 8.6 ± 3.5/ 7.3 ± 0.5 |
| mCMT971 | 9 |  | >25 | 43.2 ± 0.2/ 74.9 ± 22.5 |
| mCMR193 | 10 |  | 6.3 ± 0.4 | 15.4 ± 8.9/ 15.4 ± 4.0 |
| mCMT859 | 11 |  | >25 | 38.0 ± 17.2/ 21.8 ± 1.2 |
| mCMT949 | 12 |  | 8.7 ± 1.3 | 14.6 ± 6.9/ 12.6 ± 1.6 |
| mCMQ958 | 13 |  | 5.1 ± 2.4 | 16.3 ± 6.2/ 12.5 ± 1.0 |
| mCMQ069 | 14 |  | 3.4 ± 1.8 | 3.8 ± 1.7/ 3.3 ± 0.2 |
| mCMT467 | 15 |  | 21 ± 4.1 | 35.5 ± 14.1/ 23.7 ± 3.1 |
| mCMT954 | 16 |  | 7.6 ± 2.9 | 15.8 ± 7.0/ 13.5 ± 0.1 |

**Table 3.** Southern Aromatic rings in mCMP705.

| Compound | Entry | R | *Pf* 48 h Dd2 luc EC50 (nM) | *Pf* 72 h HEK293T/HepG2 CC50 (uM) |
| --- | --- | --- | --- | --- |
| mCMP705 | 1 |  | 1.4 ± 0.03 | 7.5 ± 3.6/ 6.8 ± 1.1 |
| mCMV869 | 2 |  | 1.1 ± 0.02 | 3.9 ± 0.1/ 3.6 ± 0 |
| mCMU303 | 3 |  | 1.1 ± 0.1 | 5.6 ± 0.2/6.5 ± 2.0 |
| mCMU291 | 4 |  | 5.6 ± 0.01 | 4.2 ± 0.9/3.7 ± 0.6 |
| mCMU290 | 5 |  | 4.1 ± 2.0 | >20/ >20 |
| mCMU135 | 6 |  | 8.7 ± 0.5 | 4.5 ± 0.2/ 3.5 ± 0.1 |
| mCMU134 | 7 |  | 1.4 ± 0.4 | 4.3 ± 0.4/4.2 ± 0.3 |
| mCMX306 | 8 |  | 11.2 ± 0.001 | >20/ >20 |
| mCMX387 | 9 |  | 26.7 ± 0.001 | >20/ >20 |
| mCMV763 | 10 |  | 10.3 ± 3.8 | >20/ >20 |
| mCMU193 | 11 |  | 3.7 ± 0.6 | >20/ >20 |
| mCMW825 | 12 |  | >30 | >12.5/ >12.5 |
| mCMW528 | 13 |  | 25.3 ± 0.008 | >12.5/ >12.5 |
| mCMZ080 | 14 |  | 14.5 ± 0.05 | >12.5/ >12.5 |
| mCMU191 | 15 |  | 7.7 ± 0.001 | >20/ >20 |
| mCMW973 | 16 |  | 2.8 ± 0.001 | 13.6 ± 0.2/ 14 ± 1.1 |
| mCMW877 | 17 |  | 14.9 ± 0.007 | >12.5/ >12.5 |
| mCMU192 | 18 |  | 15.3 ± 2.4 | >12.5/ >12.5 |

**Table 4.** Key analogs with ADME and PK profiling.

| Compound | Entry | R | *Pf* 48 h Dd2 luc EC<sub>50</sub> (nM) | *Pf* 72 h NF54 Hpx EC<sub>50</sub> (nM) | *Pf* 72 h HEK293T/HepG2 CC<sub>50</sub> (uM) | Intrinsic Hu Hep CL (uL/min/million cells) | Rat/Ms IV CL (mL/min/kg) | Hu PPB (%) |
| --- | --- | --- | --- | --- | --- | --- | --- | --- |
| KAF156 | 1 |  | 3.6 ± 2.3 | 26.7 ± 21 | 11.2 ± 6.1/9.8 ± 0.4 | 6.2 ± 1.8 | 84.3/49.2 | 94.4 |
| mCMN583 | 2 |  | 2.9 ± 0.07 | 6.9 ± 2.7 | 9.8 ± 4.0/9.5 ± 1.7 | 8.1 ± 0.3 | 93.8/27.3 | 94.7 |
| mCMQ022 | 3 |  | 10.7 ± 1.4 | ND | 47.8 ± 24.5/32.9 ± 0.4 | 6.6 ± 0.1 | 87.3/ND | 85 |
| mCMP705 | 4 |  | 1.4 ± 0.03 | 6.3 ± 2.0 | 7.5 ± 3.6/ 6.8 ± 1.1 | 5.7 ± 1.7 | 56.6/17.0 | 98.1 |
| mCMP706 | 5 |  | 4.5 ± 0.5 | ND | 23.7 ± 12.7/ 16.4 ± 2.1 | 5.1 ± 1.6 | 97/47.6 | 90.9 |
| mCMR193 | 6 |  | 6.3 ± 0.4 | ND | 15.4 ± 8.9/ 15.4 ± 4.0 | 8.1 ± 0.6 | ND | 94.8 |
| mCMQ958 | 7 |  | 5.1 ± 2.4 | ND | 16.3 ± 6.2/ 12.5 ± 1.0 | 8.0 ± 1.1 | ND | 94.5 |
| mCMQ291 | 8 |  | 4.5 ± 0.5 | ND | 8.6 ± 3.5/ 7.3 ± 0.5 | 6.3 ± 1.8 | 63.6/ND | 98.5 |
| mCMQ069 | 9 |  | 3.4 ± 1.8 | 5.6 ± 2.0 | 3.8 ± 1.7/ 3.3 ± 0.2 | 5.2 ± 0.2 | 36.3/16.3 | 99.2 |

Other trifluoro analogs had similar hepatocyte clearance values and the rat IV clearance was similar among the difluoro analogs. The key advantage of the 3,4,5-trifluoro substitution in our lead compound mCMQ069 comes from the combination of improved in vitro clearance and in vivo performance. Despite the higher protein binding (94.4% bound for KAF156 vs 99.2% for Table 4 Entry 9) this leads to a significant improvement in projected human half-life and total dose required for potential single-dose treatment and 28-day prophylaxis using the online physiologically based pharmacokinetic (PBPK) modeling from MMV. ^14^ Observation of no significant advantage over mCMQ069 upon an examination of select aromatic rings in mCMQ069 scaffold further supported the candidature of mCMQ069 (Table 5). While mCMX168 improved clearance significantly (0.4 vs 5.2; Table 5, entry 8 vs 1), it suffered a loss in potency (9.1 vs 5.6; Table 5, entry 8 vs 1).

**Table 5.**
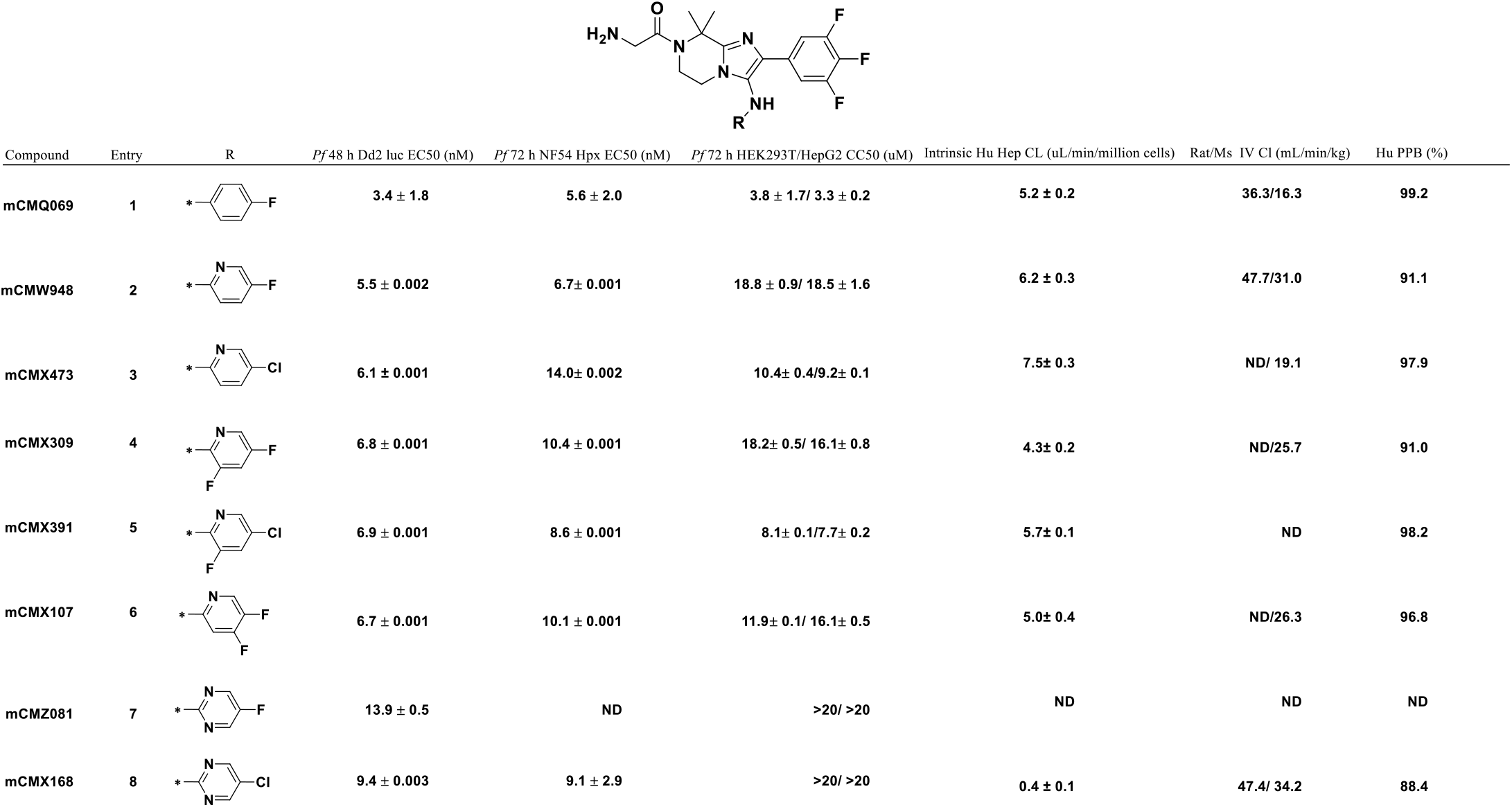
Examination of select southern aromatic rings in mCMQ069 core.

| Compound | Entry | R | *Pf* 48 h Dd2 luc EC<sub>50</sub> (nM) | *Pf* 72 h NF54 Hypx EC<sub>50</sub> (nM) | *Pf* 72 h HEK293T/HepG2 CC<sub>50</sub> (uM) | Intrinsic Hu Hep CL (uL/min/million cells) | Rat/Ms IV Cl (mL/min/kg) | Hu PPB (%) |
| --- | --- | --- | --- | --- | --- | --- | --- | --- |
| mCMQ069 | 1 |  | 3.4 ± 1.8 | 5.6 ± 2.0 | 3.8 ± 1.7/ 3.3 ± 0.2 | 5.2 ± 0.2 | 36.3/16.3 | 99.2 |
| mCMW948 | 2 |  | 5.5 ± 0.002 | 6.7 ± 0.001 | 18.8 ± 0.9/ 18.5 ± 1.6 | 6.2 ± 0.3 | 47.7/31.0 | 91.1 |
| mCMX473 | 3 |  | 6.1 ± 0.001 | 14.0 ± 0.002 | 10.4 ± 0.4/ 9.2 ± 0.1 | 7.5 ± 0.3 | ND/ 19.1 | 97.9 |
| mCMX309 | 4 |  | 6.8 ± 0.001 | 10.4 ± 0.001 | 18.2 ± 0.5/ 16.1 ± 0.8 | 4.3 ± 0.2 | ND/25.7 | 91.0 |
| mCMX391 | 5 |  | 6.9 ± 0.001 | 8.6 ± 0.001 | 8.1 ± 0.1/ 7.7 ± 0.2 | 5.7 ± 0.1 | ND | 98.2 |
| mCMX107 | 6 |  | 6.7 ± 0.001 | 10.1 ± 0.001 | 11.9 ± 0.1/ 16.1 ± 0.5 | 5.0 ± 0.4 | ND/26.3 | 96.8 |
| mCMZ081 | 7 |  | 13.9 ± 0.5 | ND | >20/ >20 | ND | ND | ND |
| mCMX168 | 8 |  | 9.4 ± 0.003 | 9.1 ± 2.9 | >20/ >20 | 0.4 ± 0.1 | 47.4/ 34.2 | 88.4 |

### *In vitro* Potency and Selectivity

mCMQ069 demonstrates activity against both liver- and blood-stage *Plasmodium* parasites. The activity against asexual blood stages has been assessed for a panel of lab-adapted *P. falciparum* strains and the mean EC_50_ values of mCMQ069 are single-digit nanomolar (e.g., 5.6 ± 2.1 nM (n=8) against *Pf* NF54 asexual blood stages employing 3H-hypoxanthine incorporation assay). Overall, regardless of assay format, whether 3H-hypoxanthine incorporation assay, 72h SYBR Green proliferation assay, or 48h viability assay with luciferase-expressing P. falciparum, we observed comparable values.

The activity against lab-adapted strains has translated to clinical isolates. Sixty *P. falciparum* field samples were tested and analyzed from June – October 2022 by IDRC laboratory in Uganda for mCMQ069 (MMV1900654) as well as 11 reference antimalarials. The test results are shown in Table S1. Reference antimalarials showed expected potency ranges against the clinical isolates (supporting information).

In addition, the *ex vivo* sensitivity of the P. falciparum and P. vivax populations from patients in Brazil, as measured at São Paulo were measured for mCMQ069 and revealed mean EC_50_ values of 30 and 3 nM, respectively (Table 6). While the team was initially concerned about the high value of 30 nM, it was determined to be an artifact of the assay where schizonticidal activity is primarily measured in the presence of serum. This difference is documented in the report and protein binding was demonstrated by assessing the activity of the lab-adapted 3D7 strain in a matrix to evaluate the potency difference in the presence of AlbuMax and serum, and between the SYBR green assay and the SMT assay. Those results showed mCMQ069 to have an EC_50_ value of 1 nM in AlbuMax-supplemented SYBR green assay and 13 nM in the presence of serum (13-fold shift). Furthermore, in the SMT assay, activities showed a further shift, yielding 63 and 130 nM in the AlbuMax- and serum-supplemented assays, respectively.

**Table 6.** *In vitro* potency and selectivity for mCMQ069.

| Assays | Activity |
| --- | --- |
| *P. falciparum* 48h Luciferase Dd2 EC<sub>50</sub> (nM) | 2.8 ± 1.9 (n=9) |
| *P. falciparum* 72h NF54 Hypoxanthine EC<sub>50</sub> (nM) | 5.6 |
| 72h NF54 HypX MPC/MIC (ng/ml) | 92/40 |
| 72h NF54 HypX Free MPC (ng/ml) | 0.7 |
| Brazilian isolates EC<sub>50</sub> (Pf, nM) | 30 |
| Brazilian isolates EC<sub>50</sub> (Pv, nM) | 3 |
| Mali isolates EC<sub>50</sub> (Po, nM) | 0.2-8.1 |
| Ghana isolates EC<sub>50</sub> (Pm, nM) | 1.96-6.6 |
| EC<sub>50</sub> (Pf, liver stage, nM) | 8 |
| indirect SMF EC<sub>50</sub> (nM) | 1.8 |
| direct SMF EC<sub>50</sub> (nM) | 300 |
| DGFA, (*Pf* male, nM) | 3 |
| DGFA, (*Pf* female, nM) | 8 |
| Cytotoxicity CC<sub>50</sub> (uM) | 3-4 |

KAF156 was included in this same assay and showed a comparable trend. Therefore, the assay format and serum binding contribute to a potency shift. KAF156 was also run in this assay and showed a highly comparable trend and value in EC_50_ potency shifts. This shift was also evident for the control compounds, artesunate and chloroquine. These data suggest that mCMQ069 is more active against asexual blood-stage *P. vivax*.

Finally, the activity of mCMQ069 was examined against clinical isolates of other species, *P. ovale* and *P. malariae* (Mali and Ghana, respectively) and provided EC_50_ ∼0.2-8.1 nM and ∼1.96-6.6 nM for *P. ovale* and *P. malariae*, respectively (Table 4). To assess the activity of mCMQ069 on liver-stage *Plasmodium* parasites, the murine parasite *Plasmodium bergheri* (*Pb*) was used as a surrogate species. mCMQ069 has a *Pb* EC_50_ = 5 nM. Subsequently, an EC_50_ of 8 nM was obtained when *Pf* liver-stage activity measured using the NF54 strain at TROPIQ. An additional request has been made for testing against *P. vivax* liver schizonts, but no other testing is expected. The team has also characterized select activity of sexual blood-stage parasites. Indirect and direct Standard Membrane Feeding Assays (SMFA; conducted by TropIQ) are the biological gold standard assessments of transmission-reducing activity (TRA). mCMQ069 resulted in an indirect SMF EC_50_ = 1.8 nM and a direct SMF EC_50_ = 300 nM. In addition, a *Pf* Dual Gamete Formation Assay (*Pf* DGFA), which evaluates male and female gamete formation, showed an EC_50_ = 3 and 8 nM for male and female gametes, respectively. These *in vitro* assays demonstrate that mCMQ069 has activity throughout the parasite life cycle, including transmission-blocking potential. The cytotoxicity assays using HEK293T and HepG2 cell lines provided CC50 in 3-4 μM range.

To assess kill kinetics against asexual blood-stage *P. falciparum*, mCMQ069 was tested in the Parasite Reduction Ratio (PRR) assay, which displayed a killing profile similar to pyrimethamine (Figure 1 and Table 7). The *in vitro* log parasite reduction ratio for mCMQ069 is 3.57 at 10 × *Pf* 3D7 EC_50_ and a parasite clearance time (PCT99.9) of ∼61 h. The killing profile included a 24-h lag phase which dictated that the speed of action be considered moderate and not fast-acting.

**Table 7.**
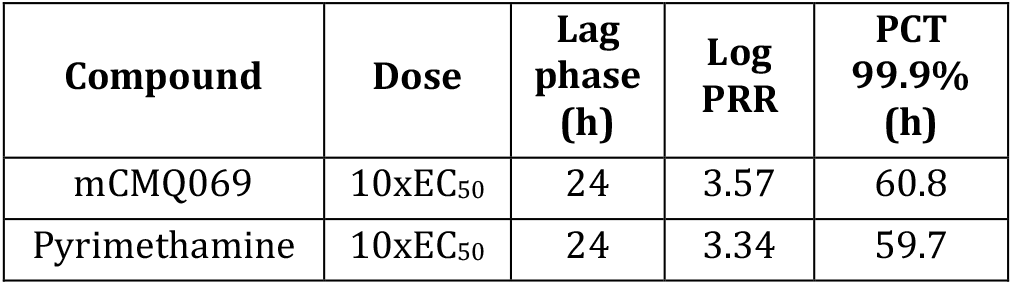
*In vitro P. falciparum* Kill Curve parameters and parasite killing rates for mCMQ069.

**Figure 1.**
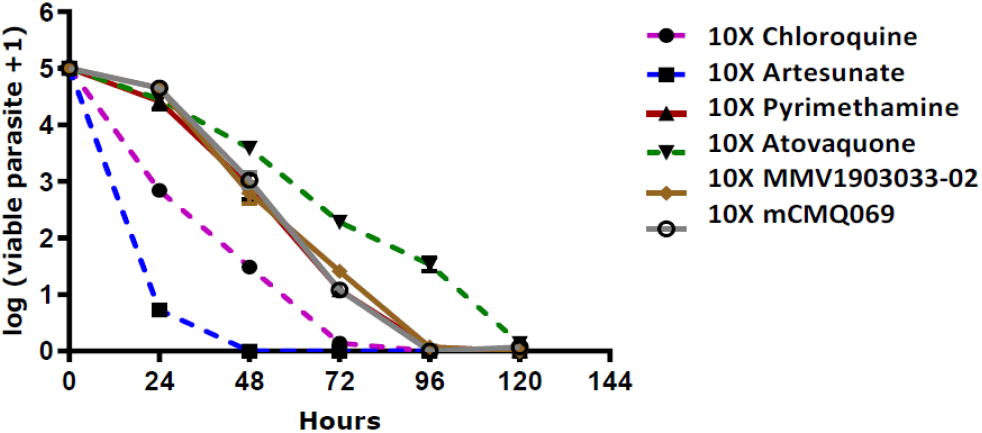
*In vitro P. falciparum* Kill Curve for mCMQ069

### *In vivo* Efficacy of mCMQ069

Oral efficacy was evaluated against erythrocytic asexual stages of *Pf* 3D7 in vivo in a humanized severe combined immunodeficient (SCID) mouse model measuring the effect on blood parasitemia. Efficacy was assessed following oral administration of the compounds as one single dose to *P. falciparum* infected NSG mice engrafted with human erythrocytes, and the effect on blood parasitemia was measured up to 20 days post-treatment. Two *Pf*SCID studies were performed at TAD to evaluate mCMQ069 for single-dose treatment. The second study included higher dose groups to increase the reliability of the analysis and to assist refinement of the human equivalent dose (HED) prediction. A summary of the parasitological response in both *Pf*SCID studies is shown in Figure 3. In total, mCMQ069 was studied at three dose levels: 10, 25, and 50 mg/kg. In both studies, the 10 mg/kg dose level reproducibly showed > 1-log reduction (>90%) in parasitemia. In the second study, a single oral dose of 25 mg/kg and above, mCMQ069 exhibited > 2 log reduction (>99%) in parasitemia at day 8 postdosing compared to untreated control mice. The average effective dose to reduce 90% parasitemia (ED_90_) was determined to be 7.0 mg/kg. The average threshold exposure required for 90% parasitemia reduction was 1,739 ng·h/mL/day (3,886 nM*h/day). In study 2, with a single dose of 10 mg/kg, 25 mg/kg, and 50 mg/kg PO, mice showed recrudescence of parasites on day 8, day 11, and day 11, respectively. These *in vivo* results support mCMQ069’s potential for a single dose cure. Significantly, by comparison, KAF156 did not reduce parasitemia after a single dose of 10 mg/kg PO and showed recrudescence of parasites on day 0. This demonstrates the differentiated efficacy profile of mCMQ069 in comparison to KAF156.

**Figure 3.**
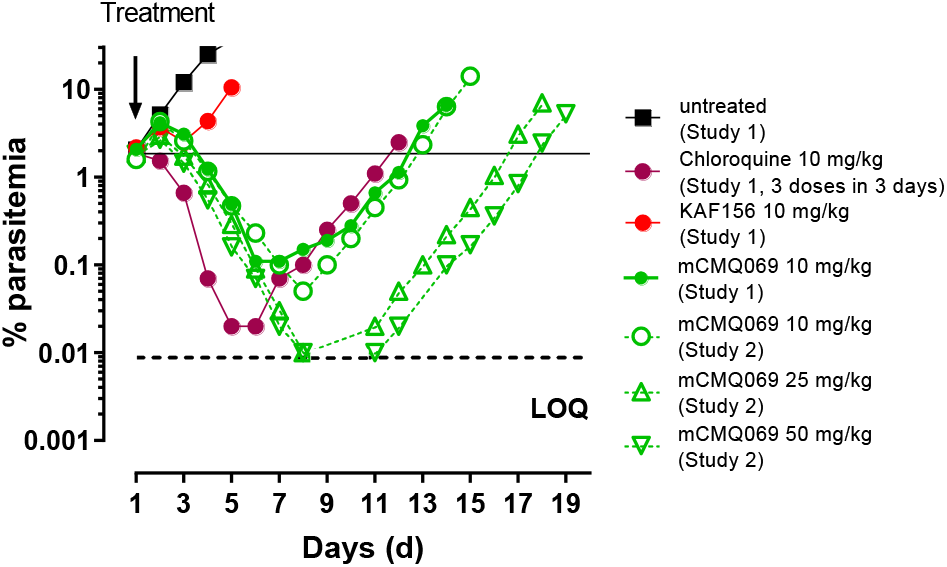
Onset of action and recrudescence for mCMQ069 and KAF156 following a single dose of 10 mg/kg (for mCMQ069 and KAF156), 25, and 50 mg/kg (for mCMQ069) on day 3 postinfection.

### *In vitro* development of resistance

Structurally, mCMQ069 is a more highly fluorinated analog of KAF156, and the presumption was that both compounds shared the same mode of action and likely had a shared mode of resistance. The studies completed to date have confirmed these assumptions and established that the mode, frequency, and degree of resistance are phenocopied.

### ADMET Properties of mCMQ069

mCMQ069 is a lipophilic base, log D at pH_7.4_ = 2.66 and pKa = 8.1, with high solubility and is classified as a DCS class II drug. The solubility in water and FaSSIF has increased when freebase was converted into a phosphate salt (Table 8). The free-base and phosphate salt of mCMQ069 are crystalline with high melting point of 201.7°C and 215°C, respectively. mCMQ069 exhibits inhibition of CYP450s, with IC50’s ranging from 0.9 to 14.8 μM for 2D6, 3A4, 2C8, and 2C9 with all others >25 μM. mCMQ069 is a highly bound compound across species in plasma protein, and microsomal and hepatocyte membranes with low in vitro microsomal and hepatocyte clearance (CL_int_).

**Table 8.** Summary of Physiochemical and ADMET Properties of mCMQ069.

| Assay | mCMQ069 free-base | Phosphate salt |
| --- | --- | --- |
| Physical form | crystalline | crystalline |
| Log P | 3.86 |  |
| Log D at pH 7.4 | 2.66 |  |
| pKa | 3, 8.14 |  |
| Melting point by DSC (°C) | 201.7 | 215 |
| Solubility in pH_3.0 citrate buffer (mg/mL) | 1.90 | >2 |
| Solubility in pH_4.7 acetate buffer (mg/mL) | >2 | >2 |
| Solubility in pH_6.8 phosphate buffer (mg/mL) | 0.50 | 1.24 |
| Solubility in pH_7.4 phosphate buffer (mg/mL) | 0.10 | 0.86 |
| Solubility in SGF (mg/mL) | >2 | >2 |
| Solubility in FaSSIF (mg/mL) | 0.14 | 0.44 |
| Solubility in FeSSIF (mg/mL) | >2 | >2 |
| Solubility in water (mg/mL) | 0.053 | >2 |
| permeability (Caco-2; Papp AB × 10<sup>6</sup> cm/s); Papp BA × 10<sup>6</sup> cm/s; efflux ratio | 0.4, 3.5, 8.8 |  |
| CYP inhibition (1A2, 2B6, 2C8, 2C9, 2C19, 2D6, 3A4 µM) | >25, >25, 2.13, 14.8, >25, 0.88, 1.3 |  |
| mouse, rat, dog, human, cyno PPB (% bound) | 98.4, 98.2, 99.3, 99.2, 98.7 |  |
| Albumax Binding (%) | 92.3 |  |
| mouse, rat, dog, human cyno microsome CL<sub>int</sub> (µL/min/mg) | 14.0, 9.8, 6.2, <9.6, NA |  |
| mouse, rat, dog, human, cyno hepatocyte CL<sub>int</sub> (µL/min/10<sup>6</sup> cells) | 28.3, 12.5, 8.4, 5.2, 11.2 |  |

### Pharmacokinetic Studies of mCMQ069

mCMQ069 freebase was dosed in vivo PO and IV to mice, rats, dogs, and cynomolgus monkeys with three animals per dosing group. Initial PK studies were done using a solution formulation with 75% PEG 400 and 25% D5W (Table 9 and Figure 4).

**Table 9.** Pharmacokinetic parameters for mCMQ069 in PO and IV mice, rat, and dogs PK studies ^a^.

|  | Mouse | Rat | Dog |
| --- | --- | --- | --- |
| Doses | 2 mpk IV, 5 mpk PO | 1 mpk IV, 5 mpk PO | 0.5 mpk IV, 2 mpk PO |
| In vivo IV clearance (mL/min/kg) | 16.3 | 36.3 | 1.9 |
| In vivo IV V<sub>ss</sub> (L/kg) | 10 | 11.9 | 9.4 |
| In vivo PO T<sub>1/2</sub> (h) | 13.4 | 5.1 | 64.5 |
| In vivo PO T<sub>max</sub> (h) | 2.2 | 3 | 8 |
| In vivo PO C<sub>max</sub> (ng/mL) | 296 | 60 | 181 |
| In vivo PO AUC (ng/mL*h) | 4,582 | 524 | 7,638 |
| In vivo PO %F | 89 | 23 | 72 |
| Blood: Plasma | 1.13 | 2.23 | 1.51 |

**Figure 4.**
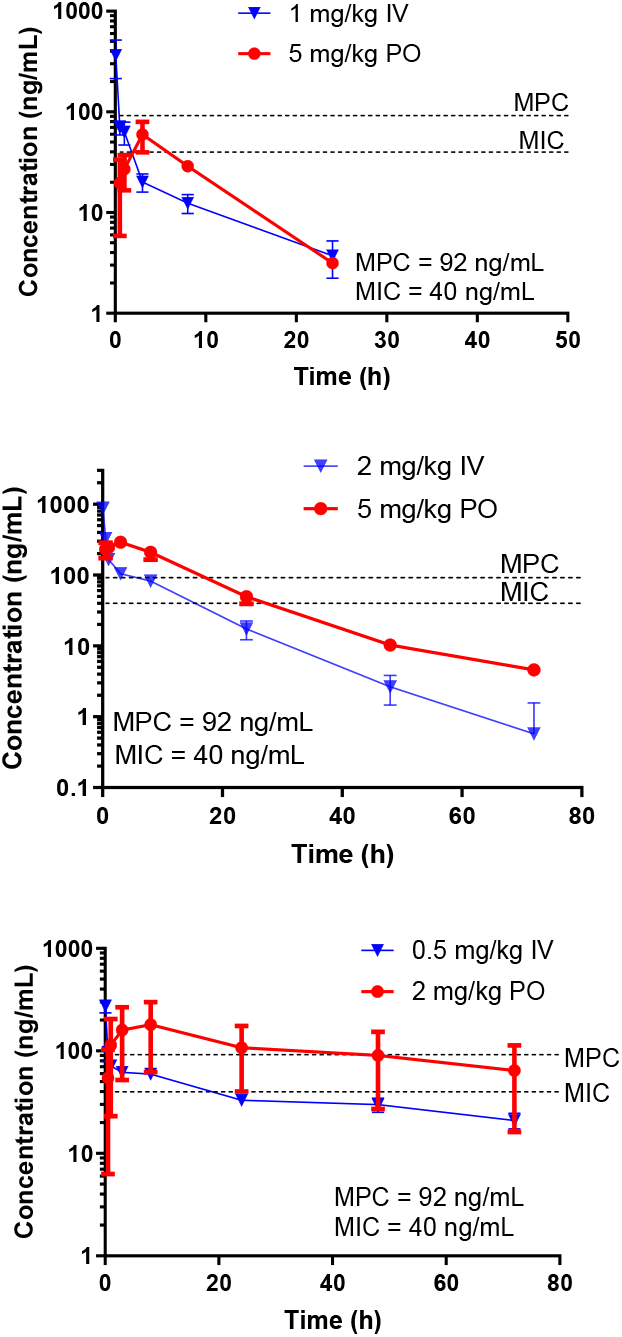
PK profile for PO and IV administered mCMQ069 in rats, mice, and dogs. The mean (±SD, n=3) plasma concentration of mCMQ069 as a function of time following the PO and IV administration.

Following a single oral dose of mCMQ069 in mice and dogs, the time to reach the maximum concentration (T_max_) was 2.2 and 8 hours, respectively, and the oral bioavailability was approximately 89% in mice and 72% in dogs, respectively. At a single 5 mg/kg oral dose of mCMQ069 in mice, mCMQ069 reached plasma exposure levels for ∼1 day over MPC (92 ng/mL) (Figure 4). On the contrary, KAF156 reached plasma levels for <0.5 days over the MPC in mice when dosed with 20 mg/kg (4x higher single dose than 5 mg/kg, not shown here). Time over MPC was confirmed in dogs as ∼2 days, supporting the longer half-life of mCMQ069 versus KAF156 in dogs (65 h vs 29 h, respectively) @ 2 mg/kg PO dosing (Figure 5A). Interestingly, under similar dosing conditions, KAF156 historical data exhibited minimal to no coverage of MPC (Figure 5B).^15^ High oral bioavailability (%F) was confirmed across species, except in rats (89% in mice, 23% in rats, 72% in dogs). Low %F in rats was most likely due to formation of additional metabolites in rats as revealed by MetID studies.

**Figure 5.**
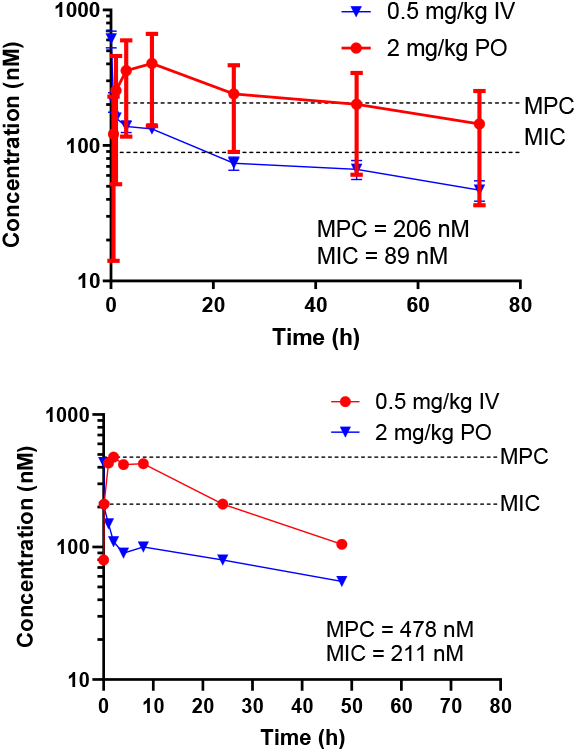
**A)** PK profile for PO and IV administered mCMQ069 in dogs. The mean (±SD, n=3) plasma concentration of mCMQ069 as a function of time following the PO and IV administration. **B)** PK profile for PO and IV administered KAF156 in dogs. The mean (±SD, n=3) plasma concentration of mCMQ069 as a function of time following the PO and IV administration. KAF156 (data has been taken from 243rd ACS Meeting San Diego, March 25, 2012^15^). MPC = Minimum Parasiticidal Concentration; MIC = Minimum Inhibitory Concentration.

These results demonstrate substantial oral absorption of mCMQ069 and slow elimination in two toxicology species.

Subsequently, mCMQ069 freebase and phosphate salt were subjected to solution and suspension dose escalation PK studies in mice and dogs to evaluate exposure and to select the final form and formulation for 7-day DRF studies (Table 10A and B).

**Table 10A.** Comparison of dose-normalized exposure in solution and suspension mice PK studies.

| API | Solution/<br>Suspension | Dose<br>(mg/kg) | Formulation<sup>a</sup> | Conc<br>(mg/mL) | d.n. AUC inf<br>((ng/mL*h)/mg/kg) | F (%) Based<br>on AUC<sub>inf</sub> |
| --- | --- | --- | --- | --- | --- | --- |
| Freebase | solution | 2 | A | 0.4 | 7209 | >100 |
| Freebase | solution | 30 | B | 6 | 6513 | >100 |
| Phosphate salt | solution | 30 | C | 6 | 7504 | >100 |
| Freebase | suspension | 30 | D | 6 | 2985 | 56 |
| Freebase | solution | 100 | B | 20 | 2554 | 48 |
| Phosphate salt | solution | 100 | C | 20 | 2414 | 45 |
| Freebase | suspension | 100 | D | 20 | 1130 | 21 |

<sup>a</sup> A = 75% PEG, 25% D5W; B = 30% PEG 400 +10% Vit E TPGS + 60%pH 4.0 Cit buffer; C = 10% PEG 400 +5% Vit E TPGS + 85%pH 4.0 Cit buffer; D = 0.5% Methyl Cellulose in pH 6.8 buffer.
AUC inf=area under the curve, initial dosing to infinity; conc=concentration; d.n. = dose-normalized; F=bioavailability.

**Table 10B.** Comparison of dose normalized exposure in solution and suspension dog PK studies.

| mCMQ069 | Solution/<br>Suspension | Dose<br>(mg/kg) | Formulation<sup>a</sup> | Conc<br>(mg/mL) | d. n. AUC inf<br>((ng/mL*h)/mg/kg) | F (%) Based<br>on AUC<sub>inf</sub> |
| --- | --- | --- | --- | --- | --- | --- |
| Freebase | solution | 5 | A | 1 | 916 | 88.9 |
| Freebase | solution | 30 | B | 3 | 408 | 39.5 |
| Phosphate salt | solution | 30 | C | 3 | 445 | 43 |
| Freebase | suspension | 33 | D | 6.6 | 633 | 61.2 |
| Freebase | solution | 100 | B | 10 | 523 | 50.7 |
| Phosphate salt | solution | 100 | C | 10 | 584 | 56.7 |
| Freebase | suspension | 110 | D | 22 | 470 | 45.5 |
| Freebase | solution | 200 | B | 20 | 598 | 58 |
| Phosphate salt | solution | 200 | C | 20 | 649 | 62.9 |
| Freebase | suspension | 330 | D | 66 | 139 | 13.4 |

<sup>a</sup> A = 75% PEG, 25% D5W; B = 30% PEG 400 +10% Vit E TPGS + 60%pH 4.0 Cit buffer; C = 10% PEG 400 +5% Vit E TPGS + 85%pH 4.0 Cit buffer; D = 0.5% Methyl Cellulose in pH 6.8 buffer.
AUC inf=area under the curve, initial dosing to infinity; conc=concentration; d.n. = dose-normalized; F=bioavailability.

Mice PK: Both freeform and salt show near dose proportional increase in AUC for solution PK. Phosphate salt shows slightly higher exposures than free base in solution at all doses ∼ 30, 100, and 300 mg/kg. Phosphate salt shows ∼1.4-fold lower dose-normalized (d.n.) AUC at 30 mg/kg in solution as compared to suspension. Phosphate salt shows ∼4.7-fold higher d.n. AUC at a higher dose of 200 mg/kg in solution as compared to 330 mg/kg in suspension. Overall, phosphate salt in solution PK is preferred due to low % PEG 400.

Dog PK: In both solution and suspension, AUC seemed to be saturated at 30 mg/kg. Phosphate salt showed slightly higher exposure in solution at ∼30 and slightly lower at 100 mg/kg than free base. Phosphate salt showed ∼2.5-fold higher d.n. AUC at 30 mg/kg in solution as compared to suspension.

Overall, phosphate salt of mCMQ069 in solution PK was selected due to low % PEG 400 for further non-GLP tox studies.

### *In vitro* Metabolism of mCMQ069

mCMQ069 was further subjected to metabolite profiling and identification in mouse, rat, rabbit, dog, monkey, and human hepatocytes. mCMQ069 was incubated with cryopreserved hepatocytes at 10 μM for up to 4 h at 37ºC followed by the analysis of the samples by LC-UV-MS. Three metabolites (M8 and M9: mono-oxygenation; M10: oxidative deamination and mono-oxygenation) were obtained across all species. In human hepatocytes, mCMQ069 and 3 metabolites (M8, M9, and M10) were detected. M8 and M9 accounting for 2.94% and 2.95% of the total drug-related components were the top two metabolites, while the M10 was only detected by MS. A select summary of the *in vitro* metabolites (10 in total) obtained from this study is provided in **Figure 6**. In mouse hepatocytes, mCMQ069 (72.1%) and 3 metabolites (M8-17.1%, M9-10.5% and M10-0.4%) were found in higher abundance than in humans. Interestingly, a total of 10 metabolites (M1-M10) with M8 and M9 major metabolites along with mCMQ069 (59.1%) were found in rat hepatocytes which could contribute to its low bioavailability in oral PK studies.

**Figure 6.**
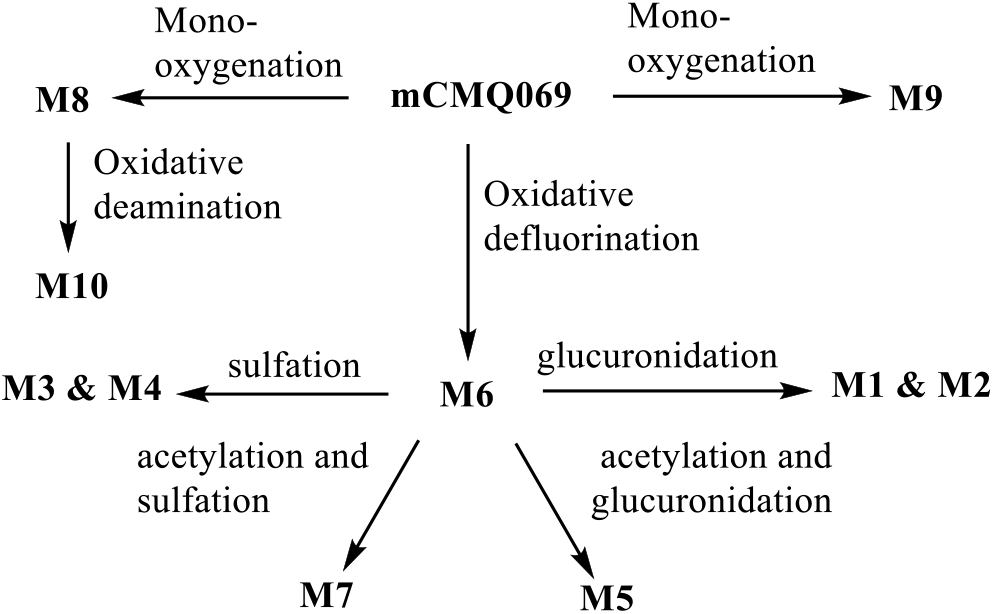
*In vitro* metabolism of mCMQ069 following incubation in mouse, rat, rabbit, dog, monkey, and human cryopreserved hepatocytes at 10 μM for up to 4 h.

### Human Dose Prediction of mCMQ069

The MMVSola tool, developed by MMV, was used to identify whether mCMQ069 has the potential as a single-dose cure for malaria. MMVSola is a mathematical human dose prediction model developed by MMV using IVIVE to assist malaria drug discovery at early and late stages.^14^ The sophisticated model combines data from preclinical *in vivo* and *in vitro* experiments including in vitro- and in vivo-derived MPC to predict the amount of drug needed to clear all malaria parasites from a patient. MMVSola predicted a 12-log kill dose of mCMQ069 to be 40–106 mg using IVIVE method. Specifically, predictions of 40 mg and 106 mg were obtained based on *in vitro* and *in vivo* data, respectively. Similarly, an IVIVE based dose range of 96–216 mg was predicted to be effective for 28- day chemoprevention. The predicted human parameters include CL = 0.31 mL/min/kg and V_ss_ = 6.8 L/kg with t_1/2_ > 200 h.

### General Synthesis and Scale-up Synthesis

The synthesis of these compounds is similar to the reported one for KAF156 in the literature. Modification of the 2-aryl ring requires an early input starting material of the appropriate isocyanide using the Groebke-Blackburn cyclization or corresponding ketone derivative (see supporting information for details). Overall yields are comparable to that of Ganaplacide and scale up costs and costs of goods assessments are in line with KAF156.

## CONCLUSIONS

We have developed a small-molecule inhibitor, mCMQ069, to improve upon the current leading antimalarial candidate, KAF156, which is currently in a Phase 3 trial likely as a 3-day treatment course of KAF156 (400 mg/day) with lumefantrine (960 mg/day). Like KAF156, mCMQ069 has a different mechanism of action than current malaria therapies, so it is a promising partner drug for existing treatments. mCMQ069 is predicted to cure an average adult malaria patient with a single, oral dose of <200 mg (TCP-1, TPP-1) with a partner drug which would protect against resistance emerging, make it more convenient and more effective at lower doses than KAF156. Preclinical modeling indicates that mCMQ069 can also serve as a single-dose malaria chemoprevention/ prophylaxis for 28 days post dose (TCP-1 & TCP-4; TPP-2), which sets it further apart from other approved malaria therapies and chemoprophylactics. mCMQ069’s high efficacy and long-acting profile has the potential to transform and simplify therapy and intervention.

## EXPERIMENTAL SECTION

### Materials and Methods

All reagents and solvents were purchased from commercial sources and used without further purification. Flash column chromatography was performed using silica gel (200−300 mesh). All reactions were monitored by TLC (pre-coated EMD silica gel 60 F254 TLC aluminum sheets and visualized with a UV lamp or appropriate stains) and/or LCMS (Waters Acquity UPLC system, 2 or 4 min run of a 10−90% mobile phase gradient of acetonitrile in water [+0.1% formic acid]). NMR spectra were obtained on Bruker AV400 or AV500 instruments, and data was analyzed using the MestReNova NMR software (Mestrelab Research S. L.). Chemical shifts (*δ*) are expressed in ppm and are internally referenced for ^1^H NMR (DMSO-d6 2.50 ppm) and ^13^C NMR (CDCl_3_ 77.16 ppm, DMSO-d_6_ 39.52 ppm).

### Chemistry

All compounds were synthesized according to the representative procedure outlined for mCMQ319 below. All compounds are >95% pure by HPLC and/or NMR analyses.

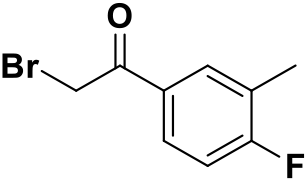

#### 2-bromo-1-(4-fluoro-3-methylphenyl)ethan-1-one

To a stirring solution of 1-(4-fluoro-3-methylphenyl)ethan-1-one (2.00 g, 13.2 mmol) in 10 mL of chloroform at room temperature was added a solution of bromine (2.54 g, 15.8 mmol, 1.2 equiv) in 15 mL of chloroform. After 15 minutes, the solution clarified and released HBr gas. The solution was allowed to stir for 15 minutes more before the solvent was removed under vacuum. The crude oil was taken to the next step without purification.

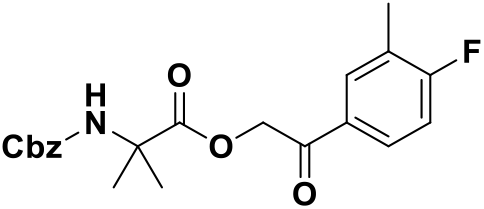

#### 2-(4-fluoro-3-methylphenyl)-2-oxoethyl 2-(((benzyloxy)carbonyl)amino)-2-methylpropanoate

The crude oil of 2-bromo-1-(4-fluoro-3-methylphenyl)ethan-1-one was dissolved in 20 mL of DMF. To the stirring solution was added solid 2-(((benzyloxy)carbonyl)amino)-2-methylpropanoic acid (3.75 g, 15.8 mmol, 1.2 equiv) and potassium carbonate (4.08 g, 29.6, 2.2 equiv) at room temperature. After 1 hour the reaction was complete by TLC and was quenched with 60 mL of water. Two extractions with 50 mL of ethyl acetate were performed and the combined organics were washed with 40 mL of water twice and an additional time with 40 mL brine. The organics were dried with sodium sulfate, filtered, and evaporated to yield 2-(4-fluoro-3-methylphenyl)-2-oxoethyl 2-(((benzyloxy)carbonyl)amino)-2-methylpropanoate as an oil (4.56 g). The material was sufficiently pure and taken to the following step without purification.

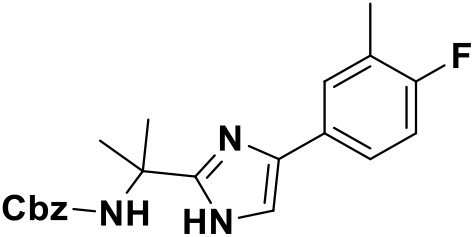

#### benzyl (2-(4-(4-fluoro-3-methylphenyl)-1H-imidazol-2-yl)propan-2-yl)carbamate

Solid ammonium acetate (10.8 g, 118 mmol, 10 equiv) was added to crude 2-(4-fluoro-3-methylphenyl)-2-oxoethyl 2-(((benzyloxy)carbonyl)amino)-2-methylpropanoate (4.56 g, 11.8 mmol) and the material was suspended in 44 mL of toluene. The flask was equipped with a reflux condenser and the mixture was brought to reflux. After 4 hours, the mixture was allowed to cool, and the solvent was removed under vacuum. To the resulting mixture of solids and oil was added 60 mL of water followed by 40 mL of ethyl acetate. The organics were separated and an additional extraction with 40 mL of ethyl acetate was performed. The combined organics were washed with 40 mL of water twice and an additional time with 40 mL of brine. The organic layer was dried with sodium sulfate, filtered, and evaporated to a red-orange oil which was purified by column chromatography utilizing a 0-100% hexane to ethyl acetate gradient. The compound eluted at 100% ethyl acetate and the fractions were evaporated to yield a colorless solid benzyl (2-(4-(4-fluoro-3-methylphenyl)-1H-imidazol-2-yl)propan-2-yl)carbamate (2.4 g, 54%).

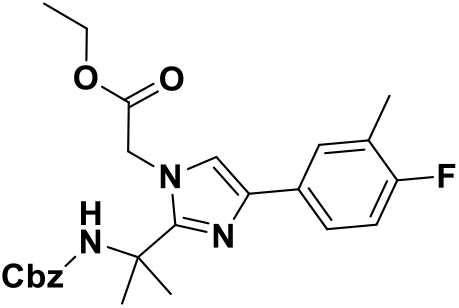

#### ethyl 2-(2-(2-(((benzyloxy)carbonyl)amino)propan-2-yl)-4-(4-fluoro-3-methylphenyl)-1H-imidazol-1-yl)acetate

Intermediate benzyl (2-(4-(4-fluoro-3-methylphenyl)-1H-imidazol-2-yl)propan-2-yl)carbamate (2.37 g, 6.4 mmol) was dissolved in 15 mL of DMF and cesium carbonate (5.3 g, 16.1 mmol, 2.5 equiv) was added to the stirring solution followed by ethyl bromoacetate (1.4 g, 8.1 mmol, 1.2 equiv) at room temperature. After 1 hour the reaction was complete by LCMS, and the reaction was quenched with 50 mL of water. The product was extracted twice with 35 mL of ethyl acetate. The combined organics were washed with 35 mL of water twice and once with 35 mL of brine. The organics were dried over sodium sulfate, filtered, and evaporated to yield crude intermediate ethyl 2-(2-(2-(((benzyloxy)carbonyl)amino)propan-2-yl)-4-(4-fluoro-3-methylphenyl)-1H-imidazol-1-yl)acetate as an oil (3.15 g).

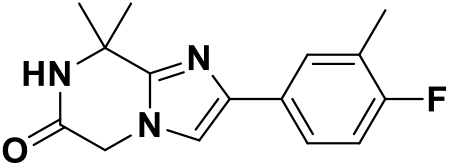

#### 2-(4-fluoro-3-methylphenyl)-8,8-dimethyl-7,8-dihydroimidazo[1,2-a]pyrazin-6(5H)-one

Intermediate ethyl 2-(2-(2-(((benzyloxy)carbonyl)amino)propan-2-yl)-4-(4-fluoro-3-methylphenyl)-1H-imidazol-1-yl)acetate (3.15 g, 6.9 mmol) was dissolved in 30 mL of methanol. A scoop of 10% w/w Pd/C was added to the solution at room temperature followed by evacuating and backfilling with hydrogen gas. The backfill process was repeated twice more. After 2.5 hours, the reaction was complete by LCMS and the mixture was filtered through a celite pad and glass frit. The celite was washed with 130 mL of ethanol and the filtrate was evaporated to yield crude 2-(4-fluoro-3-methylphenyl)-8,8-dimethyl-7,8-dihydroimidazo[1,2-a]pyrazin-6(5H)-one (1.76 g) as a solid.

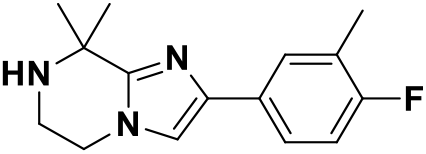

#### 2-(4-fluoro-3-methylphenyl)-8,8-dimethyl-5,6,7,8-tetrahydroimidazo[1,2-a]pyrazine

Intermediate 2-(4-fluoro-3-methylphenyl)-8,8-dimethyl-7,8-dihydroimidazo[1,2-a]pyrazin-6(5H)-one (1.76 g, 6.4 mmol) was dissolved in THF and 1.0 M borane THF complex solution (20.0 mL, 20 mmol, 3.1 equiv) was added at room temperature, and the solution was heated to reflux for 18 hours. The reaction was quenched with 20 mL of methanol and evaporated to a viscous oil. More methanol was added to quench residual borane and the solvent was once again removed. The crude yield of 2-(4-fluoro-3-methylphenyl)-8,8-dimethyl-5,6,7,8-tetrahydroimidazo[1,2-a]pyrazine (2.1 g) was much greater than the theoretical yield due to impurities from the borane and solvent but was taken onto the next step due to instability of the piperazine to silica and oxygen.

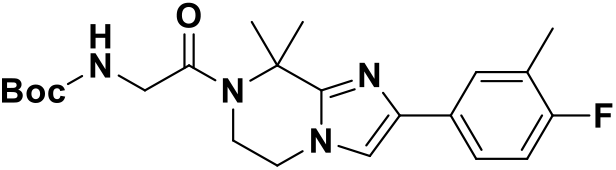

#### tert-butyl (2-(2-(4-fluoro-3-methylphenyl)-8,8-dimethyl-5,6-dihydroimidazo[1,2-a]pyrazin-7(8H)-yl)-2-oxoethyl)carbamate

Boc-glycine (2.6 g, 14.8 mmol, 2.3 equiv) and HATU (6.4 g, 11.0 mmol, 2.6 equiv) were dissolved in 7 mL of DMF at room temperature. After 10 minutes, a solution of 2-(4-fluoro-3-methylphenyl)-8,8-dimethyl-5,6,7,8-tetrahydroimidazo[1,2-a]pyrazine (2.1 g, 6.4 mmol) and DIPEA (3.5 mL, 20 mmol, 3.1 equiv) in 9 mL of DMF. The reaction was heated to 50oC for 18 hours. The reaction was quenched with 40 mL of water and the product extracted with 25 mL of ethyl acetate three times. The combined organics were washed with 25 mL of saturated sodium bicarbonate solution, 25 mL of water, and 35 mL of brine. The organics were dried over sodium sulfate, filtered, and evaporated to a crude oil which was purified by column chromatography utilizing a gradient of 0-3% MeOH/DCM. Evaporation of the column fractions yielded tert-butyl (2-(2-(4-fluoro-3-methylphenyl)-8,8-dimethyl-5,6-dihydroimidazo[1,2-a]pyrazin-7(8H)-yl)-2-oxoethyl)carbamate (2.3 g, 85%) as a white solid.

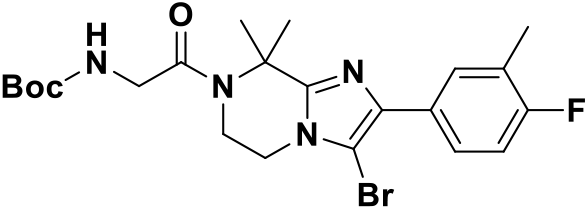

#### tert-butyl (2-(3-bromo-2-(4-fluoro-3-methylphenyl)-8,8-dimethyl-5,6-dihydroimidazo[1,2-a]pyrazin-7(8H)-yl)-2-oxoethyl)carbamate

Intermediate tert-butyl (2-(2-(4-fluoro-3-methylphenyl)-8,8-dimethyl-5,6-dihydroimidazo[1,2-a]pyrazin-7(8H)-yl)-2-oxoethyl)carbamate (0.18 g, 0.42 mmol) was dissolved in 4 mL of DCM at room temperature followed by the addition of 1.0 M bromine (0.60 mL, 0.60 mmol, 1.4 equiv) in acetic acid solution. The reation was complete within 1 hour and the product precipitated as hydrobromic acid salt. Saturated sodium bicarbonate solution (20 mL) was added to the reaction mixture and the product was extracted three times with 20 mL of ethyl acetate. The combined organics were washed with 20 mL of sodium bicarbonate solution, 20 mL of water, and 20 mL of brine. The organics were dried over sodium sulfate, filtered, and evaporated to yield crude tertbutyl (2-(3-bromo-2-(4-fluoro-3-methylphenyl)-8,8-dimethyl-5,6-dihydroimidazo[1,2-a]pyrazin-7(8H)-yl)-2-oxoethyl)carbamate as a solid (0.17 g, 79%).

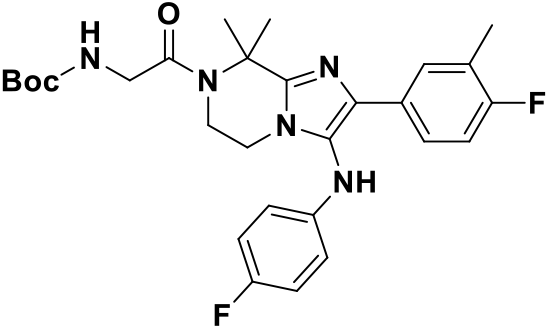

#### tert-butyl (2-(2-(4-fluoro-3-methylphenyl)-3-((4-fluorophenyl)amino)-8,8-dimethyl-5,6-dihydroimidazo[1,2-a]pyrazin-7(8H)-yl)-2-oxoethyl)carbamate

Following dissolution of tert-butyl (2-(3-bromo-2-(4-fluoro-3-methylphenyl)-8,8-dimethyl-5,6-dihydroimidazo[1,2-a]pyrazin-7(8H)-yl)-2-oxoethyl)carbamate (0.17 g, 0.35 mmol) in 4 mL of toluene, 4-fluoroaniline (0.12 g, 1.1 mmol, 3.0 equiv), xantphos (0.051 g, 0.088 mmol, 0.25 equiv), cesium carbonate (0.35 g, 1.1 mmol, 3.0 equiv), and tris(dibenzylideneacetone)dipalladium(0) (0.032 g, 0.035 mmol, 0.1 equiv) were added quickly, maintaining air-free conditions. A reflux condenser was equipped to the flask and heated to reflux for 14 hours. The reaction was quenched with 20 mL of water and extracted with 20 mL of ethyl acetate three times. The combined organics were washed with 20 mL of brine, dried over sodium sulfate, filtered, and evaporated. Column chromatography with a MeOH/DCM gradient was used to purify the product, and it eluted at 3% MeOH. Evaporation of the fractions yielded solid tert-butyl (2-(2-(4-fluoro-3-methylphenyl)-3-((4-fluorophenyl)amino)-8,8-dimethyl-5,6-dihydroimidazo[1,2-a]pyrazin-7(8H)-yl)-2-oxoethyl)carbamate (0.10 g, 54%, 85% pure as the desbrominated starting material did not separate by chromatography).

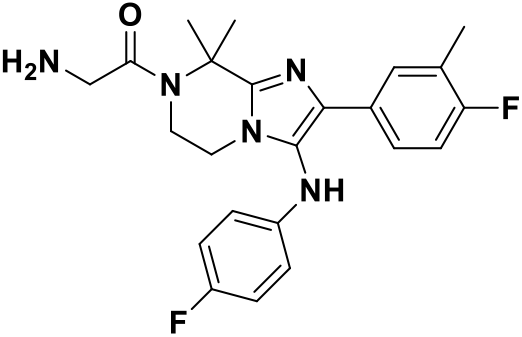

#### 2-amino-1-(2-(4-fluoro-3-methylphenyl)-3-((4-fluorophenyl)amino)-8,8-dimethyl-5,6-dihydroimidazo[1,2-a]pyrazin-7(8H)-yl)ethan-1-one

To a stirring solution of intermediate tert-butyl (2-(2-(4-fluoro-3-methylphenyl)-3-((4-fluorophenyl)amino)-8,8-dimethyl-5,6-dihydroimidazo[1,2-a]pyrazin-7(8H)-yl)-2-oxoethyl)carbamate (0.10 g, 0.19 mmol) in 2.2 mL of dioxane was added 2.2 mL 4.0 M HCl in dioxane solution (8.6 mmol, 45 equiv) at room temperature. After 2 hours, the solution was evaporated. The crude material was dissolved in 1 mL of MeOH, filtered, and purified by mass-triggered prep HPLC. The resulting TFA salt was desalted by stirring in a DCM/bicarbonate mixture. The organic layer was separated, washed with bicarbonate solution and separately with brine, dried with sodium sulfate, filtered, and evaporated. The resulting solid was lyophilized to yield a fluffy white solid 2-amino-1-(2-(4-fluoro-3-methylphenyl)-3-((4-fluorophenyl)amino)-8,8-dimethyl-5,6-dihydroimidazo[1,2-a]pyrazin-7(8H)-yl)ethan-1-one; mCMQ319 (0.023 g, 28%). LCMS: 426.4739 (M + 1) ^1^H NMR (400 MHz, DMSO-D_6_) δ 7.78 (d, *J* = 2.0 Hz, 1H), 7.67 (dd, *J* = 7.8, 2.4 HZ, 1H), 7.57 (ddd, *J* = 8.0, 5.3, 2.5 Hz, 1H), 7.09 – 6.90 (m, 3H), 6.54 (ddt, *J* = 6.6, 4.5, 2.1 HZ, 2H), 3.67 (s, 2H), 3.62 (s, 2H), 3.38 (s, 2H), 2.18 (d, *J* = 2.2 Hz, 3H), 1.84 (s, 6H).

Complete characterization of the lead molecule mCMQ069 is provided in Supporting Information.

## ASSOCIATED CONTENT

### Supporting Information

This material is available free of charge via the Internet at http://pubs.acs.org

## AUTHOR INFORMATION

### Author Contributions

The manuscript was written through contributions of all authors. All authors have given approval to the final version of the manuscript.

## ACKNOWLEDGMENTS

We would like to acknowledge Dr. Kit Bonin and Dr. Geneva Hargis for writing and graphic support.

## ABBREVIATIONS

AUC: Area under curve (exposure)
BQL: Below quantitation limit
C_last_: Final concentration measured
clogP: Calculated log partition
C_max_: Maximal concentration
C_min_: Minimum efficacious concentration
IV: Intravenous
MPC: Minimum Parasiticidal Concentration
MIC: Minimum Inhibitory Concentration
PK: Pharmacokinetic
PO: oral
SMC: Seasonal malarial chemoprevention
SPAQ: Sulfadoxine-pyrimethamine plus amodiaquine
SMT: Schizont maturation test (SMT)
TCP: Target candidate profile
T_max_: Time to peak drug concentration
TPP: Target produce profile
WHO: World Health Organization

---

Authors are required to submit a graphic entry for the Table of Contents (TOC) that, in conjunction with the manuscript title, should give the reader a representative idea of one of the following: A key structure, reaction, equation, concept, or theorem, etc., that is discussed in the manuscript. Consult the journal’s Instructions for Authors for TOC graphic specifications.

Insert Table of Contents artwork here

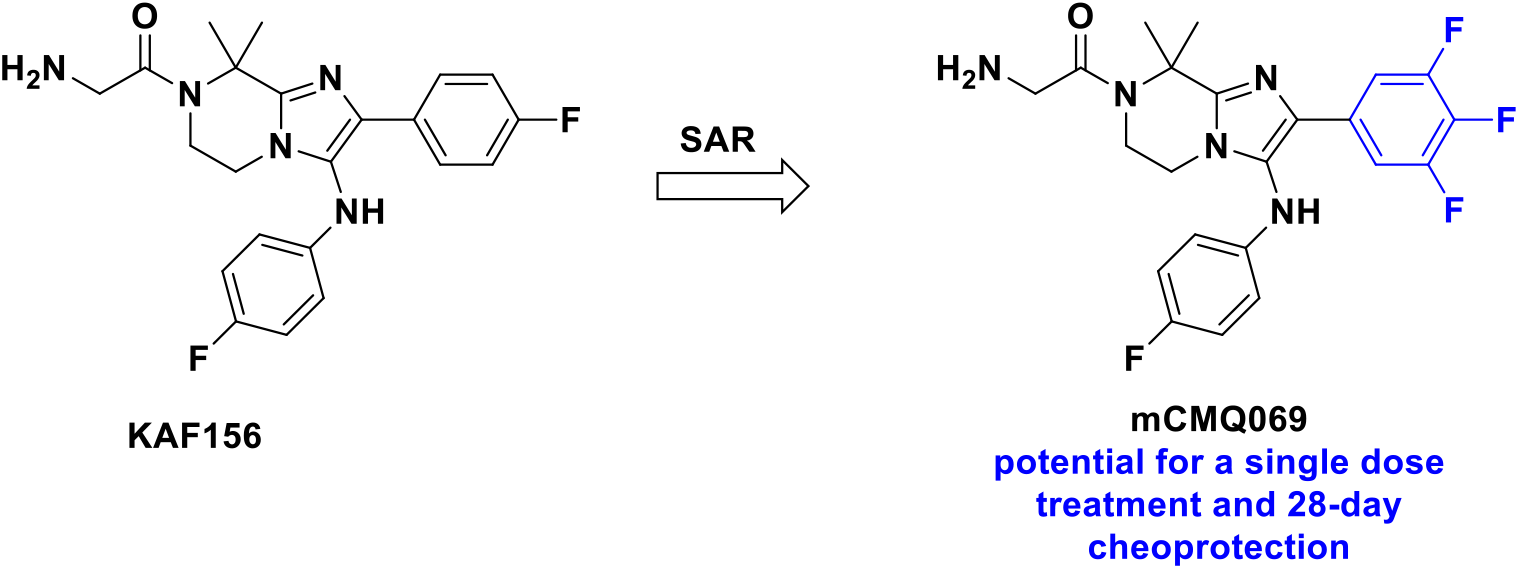

## REFERENCES

(1) World Malaria Report 2023; World Heath Organization, 2023.

(2) Malaria Reports: Malaria in the Navy and Marine Corps Active Duty Population.; Navy and Marine Corps Public Health Center, 2019.

(3) CDC. CDC Yellow Book 2024: Health Information for International Travel; 2024.

(4) Livezey, J.; Twomey, P.; Morrison, M.; Cicatelli, S.; Duncan, E. H.; Hamer, M.; Lee, C.; Hutter, J.; Mills, K.; DeLuca, J.; et al. An open label study of the safety and efficacy of a single dose of weekly chloroquine and azithromycin administered for malaria prophylaxis in healthy adults challenged with 7G8 chloroquine-resistant Plasmodium falciparum in a controlled human malaria infection model. Malaria Journal 2020, 19 (1), 336. DOI: 10.1186/s12936-020-03409-z.

(5) Saunders, D. L.; Garges, E.; Manning, J. E.; Bennett, K.; Schaffer, S.; Kosmowski, A. J.; Magill, A. J. Safety, Tolerability, and Compliance with Long-Term Antimalarial Chemoprophylaxis in American Soldiers in Afghanistan. The American Society of Tropical Medicine and Hygiene 2015, 93 (3), 584–590. DOI: 10.4269/ajtmh.15-0245.

(6) WHO recommends R21/Matrix-M vaccine for malaria prevention in updated advice on immunization. World Health Organization: 2023.

(7) Marwa, K.; Kapesa, A.; Baraka, V.; Konje, E.; Kidenya, B.; Mukonzo, J.; Kamugisha, E.; Swedberg, G. Therapeutic efficacy of artemether-lumefantrine, artesunate-amodiaquine and dihydroartemisinin-piperaquine in the treatment of uncomplicated Plasmodium falciparum malaria in Sub-Saharan Africa: A systematic review and meta-analysis. PLOS ONE 2022, 17 (3), e0264339. DOI: 10.1371/journal.pone.0264339.

(8) Hanboonkunupakarn, B.; Tarning, J.; Pukrittayakamee, S.; Chotivanich, K. Artemisinin resistance and malaria elimination: where are we now? Frontiers in Pharmacology.

(9) Sanofi. To Evaluate the Efficacy, Safety, Tolerability and Pharmacokinetics of a Single Dose Regimen of Ferroquine and Artefenomel in Adults and Children With Uncomplicated Plasmodium Falciparum Malaria (FALCI). U.S. National Library of Medicine, 2020. https://clinicaltri-als.gov/ct2/show/NCT02497612?sfpd_s=07%2F07%2F2015…sfpd_e=07%2F17%2F2015 (accessed 2022 10/27/22). Guy, R. K. Efficacy of SJ733 in Adults With Uncomplicated Plasmodium Falciparum or Vivax Malaria. U.S. Natioanl Library of Medicine, 2021. https://clinicaltri-als.gov/ct2/show/NCT04709692 (accessed 2022 10/27/22). Merck. Chemoprophylactic Activity of M5717 in PfSPZ Challenge Model. U.S. National Library of Medicine, 2022. https://clinicaltrials.gov/ct2/show/NCT04250363 (accessed 2022 10/27/22).

(10) MMV’s Pipeline of Antimalarial Drugs. Medicines for Malaria Venture, 2022. https://www.mmv.org/research-de-velopment/mmvs-pipeline-antimalarial-drugs (accessed 2022.

(11) Pharmaceuticals, N. Efficacy, Safety, Tolerability and Pharmacokinetics of KAF156 in Adult Patients With Acute, Uncomplicated Plasmodium Falciparum or Vivax Malaria Mono-infection. NCT01753323 2018.

(12) Pharmaceuticals, N. Efficacy and Safety of KAF156 in Combination With LUM-SDF in Adults and Children With Uncomplicated Plasmodium Falciparum Malaria. NCT03167242 2022.

(13) Pharmaceuticals, N. Drug-drug Interaction Study of Ganaplacide and Lumefantrine With Midazolam, Repaglinide, Dextromethorphan, Metformin, Rosuvastatin and Dolutegravir. NCT05236530 2022.

(14) https://www.mmvsola.org/ (accessed.

(15) 243rd ACS Meeting San Diego. In 2012.

